# Molecular insights into the inhibition of glutamate dehydrogenase (GDH) by the dicarboxylic acid metabolites

**DOI:** 10.1101/2021.09.28.462122

**Authors:** Barsa Kanchan Jyotshna Godsora, Prem Prakash, Narayan S. Punekar, Prasenjit Bhaumik

## Abstract

Glutamate dehydrogenase (GDH) is a salient metabolic enzyme which catalyzes the NAD^+^ - or NADP^+^-dependent reversible conversion of α-ketoglutarate (AKG) to L-glutamate; and thereby connects the carbon and nitrogen metabolism cycles in all living organisms. The function of GDH is extensively regulated by both metabolites (citrate, succinate, etc.) and non-metabolites (ATP, NADH, etc.) but sufficient molecular evidences are lacking to rationalize the inhibitory effects by the metabolites. We have expressed and purified NADP^+^-dependent *Aspergillus terreus* GDH (AtGDH) in recombinant form. Succinate, malonate, maleate, fumarate and tartrate independently inhibit the activity of AtGDH to different extents. The crystal structures of AtGDH complexed with the dicarboxylic acid metabolites and the coenzyme NADPH have been determined. Although AtGDH structures are not complexed with substrate; surprisingly, they acquire super closed conformation like previously reported for substrate and coenzyme bound catalytically competent *Aspergillus niger* GDH (AnGDH). These dicarboxylic acid metabolites partially occupy the same binding pocket as substrate; but interact with varying polar interactions and the coenzyme NADPH binds to the Domain-II of AtGDH. The low inhibition potential of tartrate as compared to other dicarboxylic acid metabolites is due to its weaker interactions of carboxylate groups with AtGDH. Our results suggest that the length of carbon skeleton and positioning of the carboxylate groups of inhibitors between two conserved lysine residues at the GDH active site might be the determinants of their inhibitory potency. Molecular details on the dicarboxylic acid metabolites bound AtGDH active site architecture presented here would be applicable to GDHs in general.

## Introduction

Glutamate dehydrogenase (GDH) is an ubiquitous metabolic enzyme which catalyses NAD^+^ - or NADP^+^-dependent reversible amination of α-ketoglutarate (AKG) to form L-glutamate in all living organisms.^1^ It, being functionally active at the branch point of the two metabolic cycles – the carbon and nitrogen cycle, is responsible for the distribution of carbon flux.^2^ GDH is being used for disease diagnosis like the blood urea nitrogen level,^3^ as malaria biomarker^4,5^ and for detection of *Clostridium difficile* infection.^6^ GDH belongs to the superfamily of amino acid dehydrogenases and the members of this group show differential substrate specificity for glutamate, phenylalanine, valine and leucine.^3^ Most of the GDHs are biologically active as hexamer (each subunit ~50 kDa) while few fungal enzymes are reported to be of tetrameric form (each subunit ~115 kDa).^7,8^ On the basis of their cofactor preference GDHs can be further categorized as NADPH, NADH or dual specific in nature.^9^ Structural studies on hexameric GDHs from several lower and higher organisms have been performed.^9–14^ In general, the monomeric subunit of hexameric GDHs has two domains: Domain-I that facilitates the substrate binding and Domain-II for NADH/NADPH binding.^10^ The mammalian GDHs are exceptional as they have an extra antenna domain which is reported to be essential in regulating the catalytic activity of the enzyme.^15^ The functional hexameric assembly of GDH is formed primarily through the interactions by domain-I of the monomers. The residues of the substrate binding pocket of the structurally characterized GDHs are highly conserved and the active site is present in the deep junction between Domain-I and Domain-II.^10^ It has been well characterized based on the structural studies that binding of coenzyme or substrate induces domain closure in GDHs. Our recent study on fungal GDH indicated formation of catalytically competent ternary complex with super closed conformation which is not observed so far for any other GDHs.^9^

The involvement of GDH in cell homeostasis has led to its tight allosteric regulation in mammalian system. In eukaryotes, the mammalian GDHs are known to be allosterically regulated by non-metabolites like GTP,^16,17^ ATP, palmitoyl CoA,^18^ NADH,^19^ steroid hormones and diethylstilbestrol (DES) inhibitors.^20^ The human GDH is associated with insulin homeostasis and its dysregulation can lead to severe diseases like genetic hypoglycemia disorder, hyperinsulinemia/ hyperammonemia syndrome (HI/HA), which occur due to loss of GTP inhibitory regulation.^21^ Some of the metabolites are also reported to control the function of mammalian GDHs. Metabolic disorders such as propionic acidemia (PAcidemia),^22^ ureagenesis and tumor growth^20,23,24^ are linked with the function of GDH. In a metabolic organic acid disorder (organic aciduria) accumulation of metabolites such as fumarate, malonate are reported.^25, 26^ Elevated levels of malonate in urine (15–70 mM) and plasma (2–10 mM) have been detected in patients suffering from malonic aciduria.^27^ Similarly in PAcidemia, maleic acid along with other precursor organic acids such as propionic acid, 3-hydroxypropionic acid and 2-methylcitric acids are also excessively excreted in urine (~5 mM).^22^ Recently, a study showed that the maleic acid specifically inhibits the kidney mitochondrial NAD^+^ linked GDH (*K*_i_ = 1.93 mM) and AKG dehydrogenase (*K*_i_ = 0.5 mM) by acting as competitive inhibitor.^22^

Additionally, there are several metabolites of TCA cycle like fructose-1, 6-diphosphate, isocitrate, fumarate and non-metabolites which are known to affect the GDH activity. Caughey and his group reported in 1957 that the 5-carbon dicarboxylate anions, AKG and glutarate act as competitive inhibitors against the beef liver GDH.^28^ The halophilic archaea *Halobacterium halobium* NAD^+^-dependent GDH is strongly inhibited by its own product AKG, glycolytic compounds (like oxaloacetate and adipate as competitive inhibitors) and the TCA cycle intermediates (like fumarate, D-glutamate, succinate, and malate as non-competitive inhibitors) in the oxidative deamination reaction.^29^ The *Clostridium botulinum* NAD^+^-dependent GDH is inhibited by the intermediates like fumarate, oxaloacetate, glutamate, glutamine and aspartate in the amination reaction.^30^ A novel large class GDH (MW ~180 kDa) in *Streptomyces clavuligerus* is also known to be inhibited by the TCA cycle intermediates such as fumarate, isocitrate, citrate, succinate, malate, oxaloacetate, glyoxylic acid and glutaric acid.^31^ Similarly, citrate an intermediate of citric acid cycle potentially inhibits the *Blastocladiella emersonii* NADP^+^-specific GDH activity.^32^ Other dehydrogenases, like succinate dehydrogenase, are also known to be inhibited by malonate, oxaloacetate and fumarate.^33^ Notably, GDH activity is affected by the metabolites of glycolytic cycle and TCA cycle and amino acids; thus existence of balanced ratio of these metabolites can render cellular homeostasis. However, none of the available reports discusses on the interactions of the metabolites with the active site of GDH. Hence, structural insights are essential to understand the molecular details of the inhibition of GDH by the carboxylate group containing metabolites.

GDH being an essential enzyme has also been a target for drug discovery against *Plasmodium falciparum*^14^ and *Arabidopsis thaliana*.^34^ Even inhibition studies on NADP^+^-dependent GDH from *Aspergillus niger* (AnGDH) led to slew of potent dicarboxylate group inhibitors such as isophthalate, 2,4-pyridinedicarboxylate and 2-methyleneglutarate which are found to be competitively more potent over monocarboxylate group inhibitors (monomethyl isophthalate, benzoic acid and butyric acid).^35^ The *in vivo* inhibitor 2-methyleneglutarate, a reaction intermediate mimic, acts as competitive inhibitor in the reductive amination of AnGDH.^36^ There is sufficient evidence of GDH inhibition by the glycolytic and TCA cycle metabolites especially during disease condition. Yet there is no direct molecular evidence of these metabolites (dicarboxylate and monocarboxylate anions) binding to GDH active site and acting as competitive or non-competitive inhibitors; hence more detailed and insightful structural studies are essential on this enzyme. In this context, glutamate dehydrogenase from *Aspergillus terreus*, an opportunistic pathogen with known industrial value for itaconic acid and livostatin production was considered. The NADP^+^-specific GDH from *A. terreus* (AtGDH) is functionally a hexameric protein with total molecular weight ~300 kDa.^37^ In this study we report high resolution crystal structures of AtGDH complexed with coenzyme NADPH and substrate mimic dicarboxylic acid metabolites like (a) malonate, (b) succinate and (c) tartrate at the active site. Our biochemical and structural studies on recombinant AtGDH reveal the interactions of few metabolites with this enzyme. The AtGDH enzyme ternary complex structures with coenzyme and substrate mimics acquire a super-closed conformation, similar to those observed for AnGDH.^9^ This study provides molecular insights into understanding of the interactions of metabolites with GDHs in general.

## Materials and methods

### Cloning, protein expression and purification of AtGDH

The *A*. *terreus* NADP^+^-specific GDH (AtGDH) cds (*gdh*A) (accession no. KF148026.1) was amplified by Q5 Polymerase using the forward primer 5′-GGATAG<u>**CATATG**</u>GAGAACCTGTACTTTCAGGGGATGTCCAACCTCCCGGTCG-3′ and reverse primer 5′-ATTG<u>**AAGCTT**</u>TTACCACCAGTCACCCTGGTCCTTCATGGCAGCAG-3′. The amplicon comprised of overhanging restriction sites at both end, and the TEV protease cleavage site (ENLYFQG) present between the *gdh*A cds and the N-terminal His_6_ tag. The construct was cloned in the pET28a(+) expression vector at the *Nde*I and *Hind*III restriction sites, followed by transformation in *E. coli* DH5α competent cells. Further, the confirmed *Atgdh* expression construct was transformed into the ΔGDH *E. coli* BL21(DE3) host cells for over-expression. The ΔGDH *E. coli* BL21(DE3) cells containing the recombinant plasmid (*Atgdh*) was inoculated into 5 ml fresh Luria Bertani (LB) broth containing kanamycin (50 μg/ml). The overnight grown culture was re-inoculated in 500 ml Luria broth and grown till the optical density O.D at 600 nm to 0.7–0.8 at 37°C. To over-express AtGDH, the cells were then induced with 0.4 mM IPTG and further grown for 12 hours with a slow shaking (80 rpm) at 22°C. The cells were harvested by centrifuging the culture at 8000 rpm and then suspended in buffer A (30 mM phosphate buffer pH 7.5). The cell suspension was lysed by ultrasonication, and then centrifuged (Multifuge, Thermo Scientific) at 4°C at 14,000*g* for 30 min. The supernatant collected was filtered through the 0.45 μm PVDF membrane (Millipore, Merck). The pre-packed Ni-NTA affinity column (GE Healthcare) equilibrated with buffer A was loaded with the filtered supernatant at a flow rate of 1 ml/min. The bound AtGDH protein was eluted with buffer B (30 mM phosphate buffer containing 150 mM imidazole, pH 7.5). N-terminal His_6_ tag of the eluted AtGDH protein was removed by TEV protease (has His_6_ tag) upon incubation for 12–16 h. The AtGDH without His_6_ tag was obtained by passing the digested sample through Ni-NTA column. Next, AtGDH sample was concentrated and further purified with the Superdex 200 pg 16/600 gel filtration column (120 ml bed volume, GE Healthcare) pre-equilibrated with buffer A. For every purification setup the purity of AtGDH was analysed using the 12% SDS-PAGE and protein quantification was done using the standard Bradford assay.

### Enzyme activity assay of AtGDH

The NADP^+^-specific glutamate dehydrogenase catalyzes the reversible reductive amination of α-ketoglutarate to L-glutamate in presence of coenzyme NADPH and excess ammonia – this is stated as forward enzyme activity. The forward activity of AtGDH was measured using the method as described earlier.^2,9,38^ The enzyme assay was performed in 1 ml reaction volume where 900 μl cocktail pH 8.0 (100 mM Tris-Cl, 10 mM α-ketoglutarate, 10 mM NH_4_Cl), a fixed volume of 0.1 mM NADPH (~0.622 *A*_340nm_) and purified enzyme (dilution required to get NADPH *A*_340nm_ 0.622 as start point) were mixed and change in absorbance at 340 nm was monitored using UV/Vis Spectrophotometer (Lambda 25, Perkin Elmer). The reaction was initiated by the final addition of NADPH. The change in absorbance (Δ*A*_340nm_) of NADPH was calculated from the initial and final time point of the reaction with linear graph (60 sec). One enzyme unit corresponds to the amount of enzyme required to oxidize/reduce one μmole of NADPH/NADP^+^ per minute under the standard assay conditions. The specific activity of enzyme was calculated and expressed as unit per mg of protein.

### Measurement of AtGDH kinetic parameters

The standard activity assay as described earlier was performed with slight modifications. The AtGDH enzyme and the substrate mimics - malonate, succinate, maleate, fumarate or tartrate was individually mixed and incubated for 10 min before reaction at room temperature. The concentration of AtGDH enzyme was kept fixed while varying the substrate mimic in the range of 0–100 mM. The pre-incubated enzyme-substrate mimic mix was directly used for the forward assay and the reaction was started by addition of NADPH at last. The relative residual activity of the AtGDH enzyme was calculated at each substrate mimic concentration by the method described above. The standard assay with no substrate mimic served as the control and referred to as *V*_o_. The *K*_i_ values were obtained by fitting the *V*_i_/*V*_o_ values to Morrison equation^39^ under the nonlinear regression (curve fit) in GraphPad Prism software (Version 8.4.2; GraphPad Prism, La Jolla, CA, USA).

### Crystallization of the AtGDH complexed with substrate mimics and coenzyme

The sitting drop vapor diffusion method was used for crystallization of AtGDH at 22°C. Several commercially available crystallization screens such as JCSG-plus (Molecular Dimensions), PEG suite (Qiagen), PEG Rx (Hampton Research), PEG/Ion (Hampton Research) and Index (Hampton Research) were used to get the initial crystal hits. The AtGDH binary complex was obtained by mixing the pure AtGDH protein (20 mg/ml) and the coenzyme NADPH till final concentration 6 mM and then pre-incubation for 30 min at ice cold condition. The crystallization drops containing AtGDH-NADPH binary complex and mother liquor with 1:1 ratio were set up in the screening tray using the automated Phoenix robot (Art Robins) at Protein Crystallography Facility, Indian Institute of Technology Bombay. The crystallization trays were incubated at 22°C in vibration free incubator. It was surprising to observe that most of the crystal hits appeared in the crystallization conditions containing dicarboxylate anions (succinate, malonate, tartrate and citrate). The crystals of tartrate (TLA) bound to AtGDH-NADPH binary complex, hereafter designated as AtGDH-TLA-NADPH appeared in the 0.2 M sodium tartrate dibasic, 20% PEG3350 pH 7.3 and those grew to the maximum size in five weeks.

The AtGDH ternary complex was prepared by mixing the AtGDH (12 mg/ml) and the coenzyme NADPH (10 mM) and substrate α-ketoglutarate (50 mM) for 10 min at 22°C. The hanging drop method was used to set up the drops with 1:1 ratio of mother liquor to AtGDH ternary complex. The crystals appeared in the 0.8 M succinic acid (SIN), pH 7.0 after 5 days of incubation at 22°C. Later, on solving the structure it was identified that the substrate α-ketoglutarate was replaced by the succinic acid to form the AtGDH-succinate-NADPH crystals which is designated as AtGDH-SIN-NADPH. Similarly, the substrate α-ketoglutarate was also identified to be replaced by malonate. The crystals of malonate (MLI) bound AtGDH-NADPH complex is designated as AtGDH-MLI-NADPH which appeared in 2.4 M sodium malonate pH 7.0 after one week of incubation.

### X-ray diffraction data collection and processing

All the diffraction data sets were collected from the frozen crystals by the rotation method. The crystals of AtGDH-TLA-NADPH, AtGDH-SIN-NADPH and AtGDH-MLI-NADPH individually, were harvested using nylon cryo-loop from the drop and transferred to their respective cryoprotectants (mother liquor with 30% glycerol) and then flash-frozen to liquid nitrogen stream at 100 K. The diffraction data were collected by rotating method at the home source using the Cu*Kα* X-ray radiation which is generated by the Rigaku Micromax 007HF X-ray generator fitted with a Rigaku R-Axis IV++ detector (Protein Crystallography Facility, IIT Bombay). All the diffraction data sets were processed (indexing, integration and scaling) using XDS software,^40^ following which the recorded intensities were converted to structure factor by the F2MTZ and CAD program of CCP4.^41^ The data collection statistics are presented in Table-1.

**Table 1:** Data collection and refinement statistics

| Data collection<sup>a</sup> | AtGDH-TLA-NADPH | AtGDH-SIN-NADPH<sup>#</sup> | AtGDH-MLI-NADPH |
| --- | --- | --- | --- |
| Space group | *I*222 | *I*222 | *I*222 |
| Unit cell parameters<br>*a*, *b*, *c* (Å)<br>*α*, *β*, *γ* (°) | 119.86, 153.09, 259.27<br>90.0, 90.0, 90.0 | 119.88, 153.75, 259.95<br>90.0, 90.0, 90.0 | 120.15, 153.74, 259.88<br>90.0, 90.0, 90.0 |
| Resolution (Å) | 39.29 – 2.90 (3.0 – 2.90) | 38.68 – 1.73 (1.83 – 1.73) | 39.39 – 1.74 (1.84 – 1.74) |
| Wavelength (Å) | 1.5418 | 1.5418 | 1.5418 |
| Temperature (K) | 100 | 100 | 100 |
| Observed reflections | 356102 (34113) | 1750773 (214326) | 1758739 (242177) |
| Unique reflections | 53081 (5072) | 246054 (36141) | 244015 (37079) |
| Completeness (%) | 99.8 (100.0) | 99.2 (94.7) | 99.8(99.1) |
| *R*<sub>merge</sub> (%) | 42.6 (159.2) | 11.0 (91.5) | 10.3 (83.2) |
| *R*<sub>meas</sub> (%) | 46.2 (168.9) | 11.9 (100.1) | 11.1 (90.4) |
| *I*/*σ I* | 4.8 (1.41) | 15.38 (2.10) | 16.72 (2.39) |
| *CC*<sub>1/2</sub> (%) | 96.6 (63.6) | 99.8 (72.2) | 99.9 (76.6) |
| Redundancy | 6.7 (6.72) | 7.11 (5.93) | 7.2 (6.53) |
| **Refinement** |  |  |  |
| Resolution (Å) | 39.29 – 2.9 | 38.68 – 1.73 | 39.39 – 1.74 |
| No. of reflections (working set/<br>test set) | 50428/ 2655 | 233751/ 12303 | 231826/ 12202 |
| *R*<sub>factor</sub> (%) | 22.2 | 14.6 | 13.9 |
| *R*<sub>free</sub> (%) | 26.9 | 16.7 | 16.3 |
| **Number of atoms** |  |  |  |
| Protein | 10371 | 10574 | 10576 |
| Water | 87 | 1608 | 1857 |
| Tartrate | 30 | 0 | 0 |
| Malonate | 0 | 0 | 70 |
| Succinate | 0 | 24 | 0 |
| NADPH | 144 | 0 | 144 |
| ADP-ribose | 0 | 120 | 0 |
| **Average isotropic B- factor for active site ligands(Å<sup>2</sup>)** |  |  |  |
| Tartrate | 49.3 | 0 | 0 |
| Malonate | 0 | 0 | 27.8 |
| Succinate | 0 | 17.4 | 0 |
| NADPH | 36.8 | 0 | 17.5 |
| ADP-ribose | 0 | 17.4 | 0 |
| **Average isotropic B- factor (Å<sup>2</sup>)**<br>**of all atoms** | 41.0 | 19.8 | 20.45 |
| **R.M.S.D** |  |  |  |
| Bond distance (Å) | 0.01 | 0.023 | 0.024 |
| Bond angle (°) | 1.5 | 2.10 | 2.28 |
| **Protein geometry** |  |  |  |
| Ramachandran plot favoured (%) | 89.7 | 92.1 | 92.1 |
| Ramachandran plot allowed (%) | 10.1 | 7.4 | 7.4 |
| Ramachandran plot outliers (%) | 0.2 | 0.5 | 0.5 |
| **PDB ID** | 7ECT | 7ECR | 7ECS |

<sup>a</sup>Values in parentheses correspond to highest resolution shell. AtGDH-TLA-NADPH: AtGDH structure complexed with tartrate (TLA) and NADPH, AtGDH-SIN-NADPH<sup>#</sup>: AtGDH structure complexed with succinate (SIN) and NADPH, AtGDH-MLI-NADPH: AtGDH structure complexed with malonate (MLI) and NADPH. <sup>#</sup>In this structure nicotinamide ring of NADPH is removed.

### Structure determination and refinement

The molecular replacement (MR) method was used to obtain the initial phases. Matthews coefficient (*V*_M_)^42^ calculated for the AtGDH-SIN-NADPH crystals was 4.05 Å^3^ Da^−1^ that corresponds to the presence of three AtGDH molecules in the asymmetric unit with 69.6% solvent content. Further the MR module of PHASER^43^ was run to find the orientations of AtGDH molecules using the AnGDH ternary complex (5XVX) as search model as latter shares 88% sequence identity with former protein. PHASER run placed three template models in the correct orientation. The AtGDH-SIN-NADPH structure was further manually built using COOT^44^ and refined using REFMAC5.^45^ Once protein part of the AtGDH-SIN-NADPH structure was built, visual inspection of the sigma-A weighted *F*_*o*_-*F*_*c*_ omit map indicated presence of succinate and NADPH without nicotinamide group in the active site although α-ketoglutarate (AKG) and NADPH were used for co-crystallization of the ternary complex. Subsequently, the ligands with correct orientation were modeled in the electron density and the complexed structure was further refined for few cycles in REFMAC5. Next, the unresolved electron density peaks above 3σ level in the sigma-A weighted *F*_*0*_-*F*_*c*_ were satisfied upon adding the solvent molecules. The structure was further refined with multiple rounds of refinement cycles until all positive peaks in the electron density were satisfactory and acceptable *R*_factor_ and *R*_free_ were obtained.

The structures of AtGDH-MLI-NADPH and AtGDH-TLA-NADPH were determined using the coordinates of subunit A of AtGDH-SIN-NADPH structure as a template. The Matthews coefficient (*V*_M_) calculated for AtGDH-MLI-NADPH and AtGDH-TLA-NADPH crystals were 4.04 and 4.03 Å^3^ Da^−1^, respectively with three AtGDH molecules in the asymmetric unit. After the refinement of the protein molecules, the electron density in the active sites of AtGDH-MLI-NADPH and AtGDH-TLA-NADPH structures indicated presence of malonate and tartrate, respectively along with cofactor NADPH. Although AtGDH-AKG-NADPH ternary complex was set for crystallization but the malonate from the mother liquor replaced the actual substrate AKG in the structure. Similarly, the AtGDH-NADPH binary complex was set for crystallization but tartrate from the crystallization mother liquor got bound to the active sites. Next, the ligands were correctly built inside the sigma-A weighted *F*_*o*_-*F*_*c*_ positive omit map at the active site pocket. Careful inspection and manual building was done in COOT and refinement in REFMAC5 as mentioned above. The refinement statistics of these structures are presented in Table-1. The stereochemistry of the residues in all three built structures was analyzed by Ramachandran plot using PROCHECK.^46^ The PyMol Molecular Graphics System, Version 2.1.1, Schrödinger, LLC and UCSF Chimera 1.13 (http://www.rbvi.ucsf.edu/chimera) were used to prepare the structural figures.

## Results

### Enzyme activity and inhibition of AtGDH

*A. terreus* glutamate dehydrogenase (AtGDH) was expressed successfully in soluble form as N-terminal His_6_ tagged protein. The His_6_ tag was efficiently cleaved by TEV protease and AtGDH was purified to almost 95% purity. AtGDH follows Michaelis-Menten kinetics and is functionally active as a hexamer. The specific activity of this pure enzyme for conversion of α-ketoglutarate to L-glutamate (forward reaction) was observed to be 160 U/mg.

The inhibition effects of AtGDH forward reductive amination reaction by malonate, succinate, maleate, fumarate and tartrate were measured (Figure 1). It is also observed that 20 mM of malonate and fumarate individually caused ~40% and 35% decrease in the enzyme activity, respectively. Similarly, when succinate and maleate were tested separately, both caused about ~30% decrease at 20 mM concentration. Low concentrations (till 20 mM) of tartrate resulted increase in the enzyme activity, but the metabolite caused ~15% inhibition at 50 mM and above (Figure 1A). The calculated *K*_i_ values are 19.5, 23.2, 36.2, 39.1 and 50.8 mM for malonate, fumarate, maleate, succinate and tartrate, respectively (Figure 1B). These data clearly indicate that the dicarboxylic acid group containing metabolites are able to inhibit the AtGDH enzymatic activity at higher concentrations. However, the inhibition potencies of these metabolites towards AtGDH are much lower as compared to isophthalate^36^ which is a well-known highly potent inhibitor of GDHs (Figure 1B). Molecular basis of the interactions of these metabolites with GDH are discussed later.

**Figure 1.**
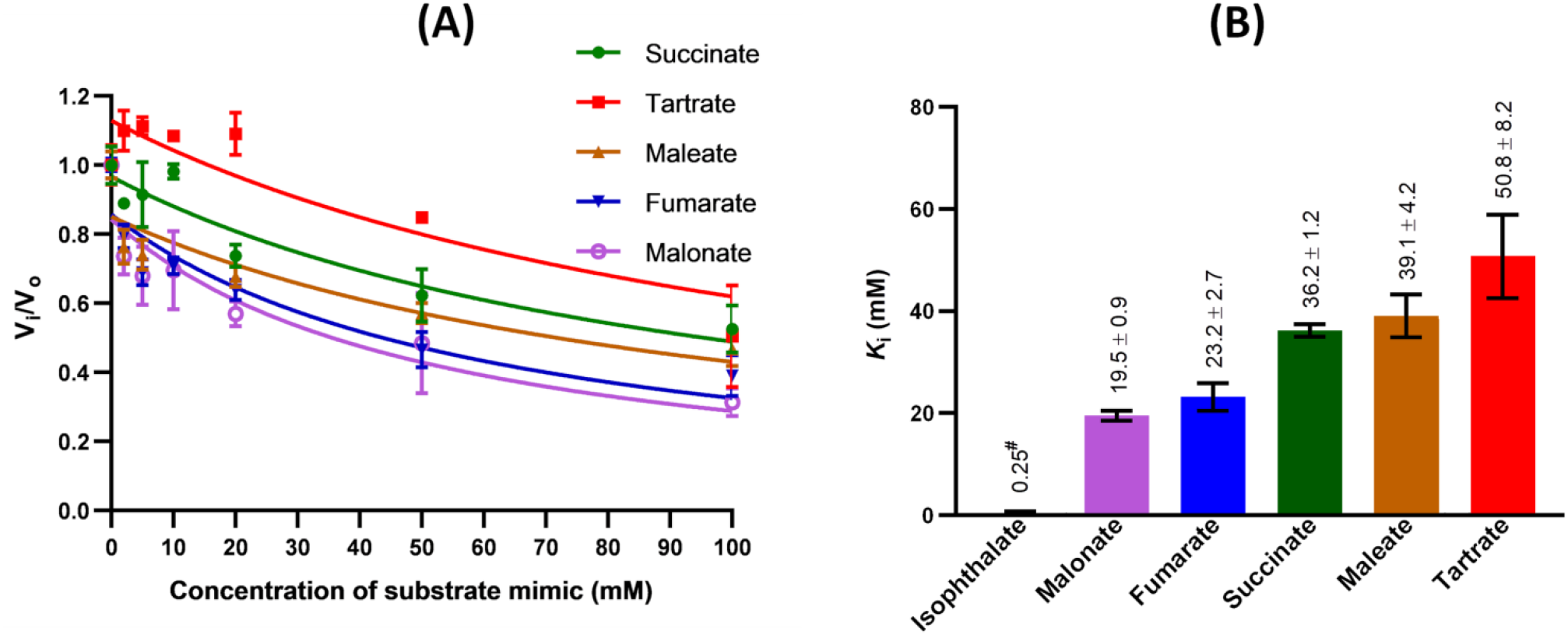
*A. terreus* GDH (AtGDH) inhibition by dicarboxylic acids. (A) Inhibition kinetics of AtGDH in presence of the five substrate mimics namely, tartrate (red line, 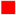), maleate (brown line, 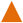), succinate (green line, 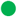), fumarate (blue line, 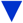) and malonate (pink line, 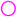). (B) Bar graph represents the *K*_i_ values of substrate mimics and inhibitor isophthalate against the forward activity of AtGDH. The experimental *K*_i_ values of substrate mimics and the reported *K*_i_ value of isophthalate^36^ (marked in #) are labelled. The experiments are done in triplicate (n = 3), and the error bar represents the standard deviation.

### Structural fold of AtGDH

The structures of AtGDH bound to coenzyme NADPH and substrate mimics- succinate, malonate and tartrate have been determined at 1.73, 1.74 and 2.9 Å resolution, respectively. The succinate, malonate and tartrate bound AtGDH structures are referred as AtGDH-SIN-NADPH, AtGDH-MLI-NADPH and AtGDH-TLA-NADPH, respectively. All these AtGDH complexed crystals belong to the same space group *I*222 with similar unit cell parameters and contain three protein molecules in the asymmetric unit. However, the crystallographic symmetry operation generates the functional hexameric biological assembly (Figure 2A). The structures reported here have been refined appropriately with good refinement statistics (Table 1).

**Figure 2.**
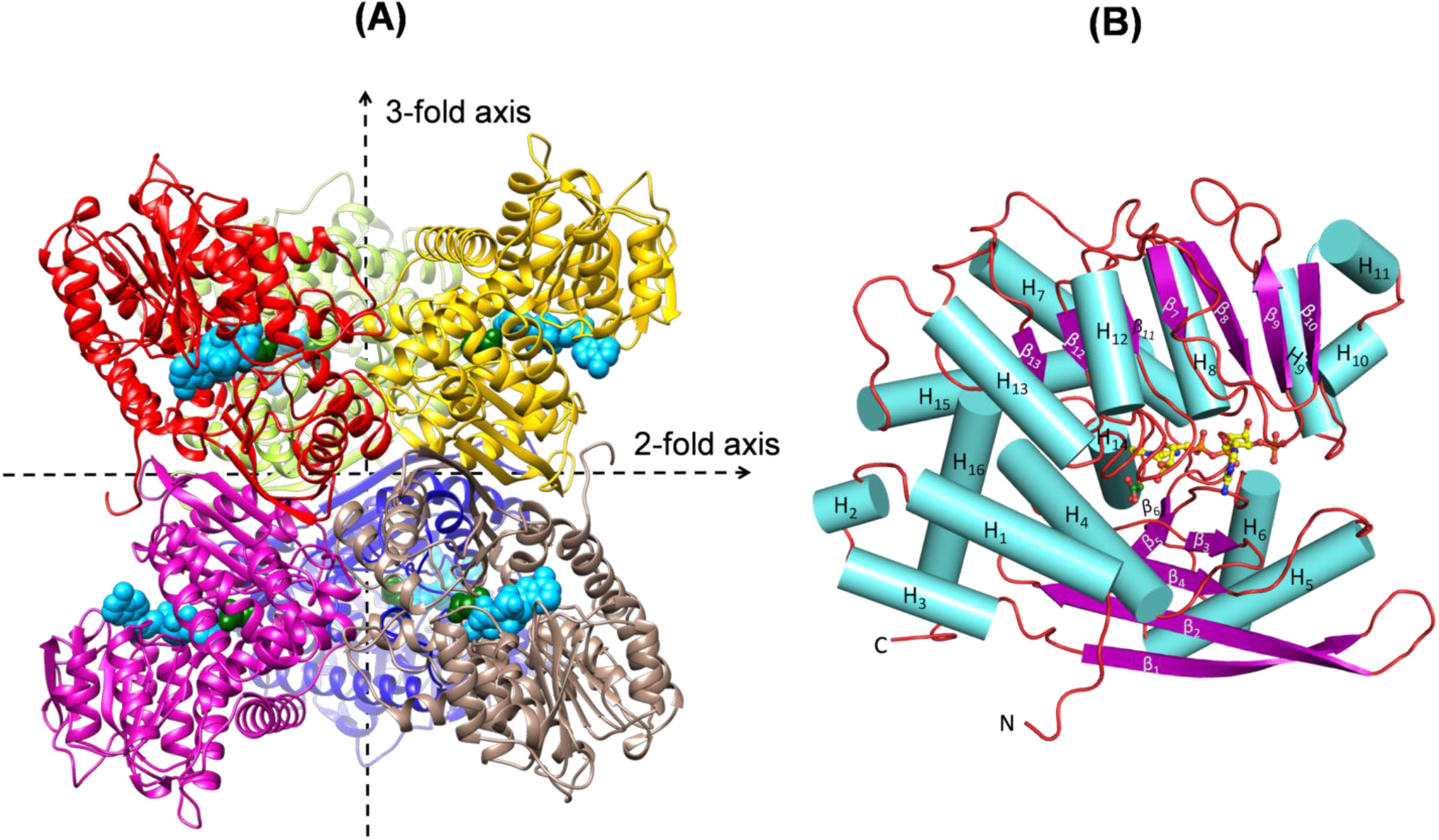
Structural fold of AtGDH. (A) Cartoon representation of hexameric AtGDH structure. The monomers are shown in different colors. Crystallographic axes are shown as dotted arrows. The bound malonate and NADPH are shown as green and cyan spheres, respectively. (B) AtGDH monomer is shown as cartoon with α-helices marked as ‘H’, β-strands marked as ‘β’ with their numbers and the loops in brick red color. The malonate (dark green) and the coenzyme NADPH (yellow) bound to the active site are shown as ball and stick models.

The structures of the monomers of AtGDH in three different complexes reported here are almost identical. A monomer of AtGDH-MLI-NADPH trimeric structure was aligned to the equivalent monomers of AtGDH-SIN-NADPH and AtGDH-TLA-NADPH using Secondary Structure Matching (SSM) superposition which produced average root mean square deviation (r.m.s.d.) values of ~0.15 Å and 0.27 Å, respectively. The three subunits of the individual structure also superimpose very well with each other (average r.m.s.d. ~0.26 Å), indicating no structural differences among the monomers. The structures of AtGDH bound to NADPH and the substrate mimics (malonate/ tartrate/ succinate) are almost identical and therefore the A-subunit of AtGDH-MLI-NADPH structure has been used here to describe the overall structural fold of this fungal enzyme.

The structure of AtGDH comprises of 16 α-helices and 13 β strands numbered as H_1_-H_16_ and β_1_-β_13_, respectively (Figure 2B); similar to those reported in *A*. *niger* GDH (AnGDH) structure (5XVX).^9^ Each subunit of AtGDH has two domains (I and II); Domain-I ranges from residues 1–190 and the last portion of the C-terminal (residues 436–460). Domain-I is involved in substrate recognition and subunit-subunit interactions, facilitating the formation of hexameric assembly. Domain-II comprises of residues 191–435 and has modified Rossmann like fold (αβα) for the dinucleotide binding. These two domains are linked by hinge like pivot helix (H_15_) which helps in Domain-II movement for the catalytic reaction. It is quite well known that the three dimensional structural fold of monomers of hexameric GDHs are well conserved in bacteria, fungi and plants. The overall tertiary structural fold of AtGDH is almost identical to its closer orthologous enzyme AnGDH (5XVX) as alignment of the monomers of these two enzymes produced r.m.s.d value of 0.34 Å (SSM superposition for 455 Cα atoms). AtGDH structure is also similar to the structural fold of *Corynebacterium glutamicum* GDH (CgGDH) (5IJZ), *Clostridium symbiosum* GDH (CsGDH) (1BGV) and *Plasmodium falciparum* GDH (PfGDH) (2BMA). The hexameric structure of AtGDH (Figure 2A) can be represented as ‘dimer of trimers’ or ‘trimer of dimers’ similar to that observed in CsGDH^7^ as well as in AnGDH.^9^ The hexamer is formed by several inter dimeric and trimeric interactions. The inter-trimeric interactions are formed by the residues in β_3_ (72–77), loop (146–148), β_5_ (163–170), loop (174– 177), β_6_ (186–188), H_14_ (390–398) and H_16_ (454–459) regions. The inter-dimeric interactions are mainly formed by the Domain-I residues: N-terminal initial residues, β_1_ (38–53), β_2_ (60–66), H_5_ (139–143) and the C-terminal end residues. These dimeric and trimeric interface interactions are mostly conserved in homologous AnGDH as well.^9^

Notably, the subunits in AtGDH-MLI-NADPH, AtGDH-SIN-NADPH and AtGDH-TLA-NADPH structures acquire super closed conformation similar to that reported in the catalytically competent ternary complex of AnGDH bound to substrate α-ketoglutarate and coenzyme NADPH (5XVX) (Figure 3A). The mouth opening, as measured by the distance between the Cα atom of the cleft residues Lys122 (Domain-I) and Arg282 (Domain-II), was found to be around ~6 Å (Figure 3A, inset). It is surprising to find that AtGDH structures adopt the super closed conformation even though the actual substrate is not bound to the active site of the enzyme. Despite having almost identical structural fold, AtGDH structure has subtle differences at the regions of amino acid insertion- loop 1 (262–263) and deletion- loop 2 (294–295) (Figure 3B, 3C) as compared to AnGDH. Implication of these structural differences on the functional properties of AtGDH needs further investigation.

**Figure 3.**
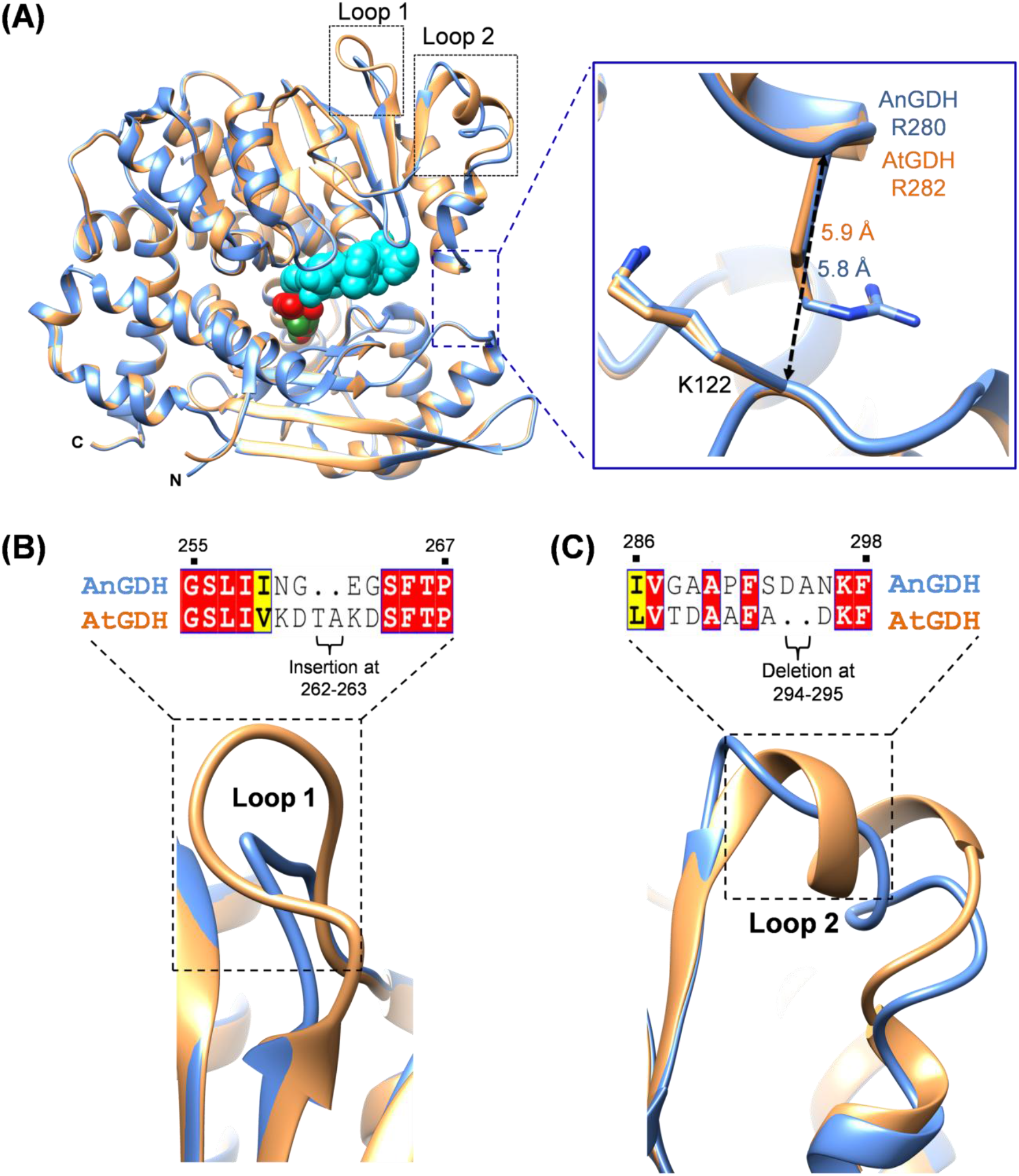
Comparison between the ligand bound AnGDH (blue carbon) and AtGDH (orange carbon) structures. (A) The structure of AnGDH (blue) ternary complex (5XVX) bound with α-ketoglutarate (red sphere) and NADPH (cyan sphere) is superposed to the AtGDH (orange) structure bound to malonate (green sphere) and NADPH (cyan sphere). The inset shows the zoom in view of the mouth opening measured by the distance between identical position arginine and lysine residues (sticks) in the structures. (B) Depiction of the region of amino acid insertion (262-263) in Loop 1 of AtGDH as compared to AnGDH. (C) The conformational differences in Loop 2 regions of AtGDH and AnGDH due to the deletion of two residues (294-295) in the former enzyme.

### Interactions of coenzyme NADPH with the active site of AtGDH

The sigma-A weighted *F*_*o*_-*F*_*c*_ omit maps of AtGDH-MLI-NADPH and AtGDH-TLA-NADPH structures confirm the presence of bound coenzyme NADPH at the active site of all the subunits. Figure 4A represents the *F*_*o*_-*F*_*c*_ omit electron density indicating presence of NADPH and malonate in the AtGDH-MLI-NADPH structure. Since all subunits of these structures are nearly identical, the interactions of the coenzyme (NADPH) with the enzyme are described here with respect to subunit A of AtGDH-MLI-NADPH structure. NADPH binds in an extended conformation at the active site of AtGDH with its reactive nicotinamide moiety buried deep inside the cavity. The coenzyme is primarily held in the binding pocket (Figure 4B, 4C) by several interacting residues from Domain-II, while only two residues from Domain-I are involved in this interaction; earlier similar results were reported for AnGDH as well.^9^ NADPH binding pocket of AtGDH is highly positively charged (Figure S1). The oxygen atom in the amide group of nicotinamide ring forms a hydrogen bond with Thr197 and an electrostatic interaction with Arg193. The ribose moiety of nicotinamide occupies the cavity formed by polar residues. The 2′-OH group of ribose forms hydrogen bonding interaction with the side chains of Arg82, Asp154 and Asn346. The 3′-OH group of ribose is stabilized by the backbone amide of Asn346 and interacts with Gln322 *via* water. The pyrophosphate group of NADPH binds at the stretch of nucleotide binding motif (GXGXXA) ranging from residues 228–233, by interacting with the backbone. The hydroxyl group on pyrophosphate is largely stabilized by the surrounding water molecules. The 3′-OH group of adenosine ribose is stabilized by the Ser229. Like AnGDH, the Asp252 in AtGDH structure flips away and interacts with Lys279 and Gln284 in order to accommodate the 2′-phosphate group. The 2′-phosphate group of adenosine ribose occupies the densely positively charged region aligned by Lys122, Ser253, Lys279 and Gln284 and forms direct side chain interactions. The adenosine nucleotide lies exposed and its hydroxyl group is stabilized by the residues His84, Ile155 and Thr321 (Figure 4B, 4C). Surprisingly, the nicotinamide ring is cleaved off from NADPH to form ADP-ribose (ADP-ribose) in the AtGDH-SIN-NADPH structure as indicated by clear *F*_*o*_-*F*_*c*_ omit map (Figure 4D); but the remaining portion of the coenzyme is intact and binds in the similar manner as described above. A striking difference is that 1′-OH group of nicotinamide ribose is interacting with Asn379 side chain which has adopted an alternate conformation. However, how the nicotinamide ring of NADPH got eliminated during the crystallization process remains unclear.

**Figure 4.**
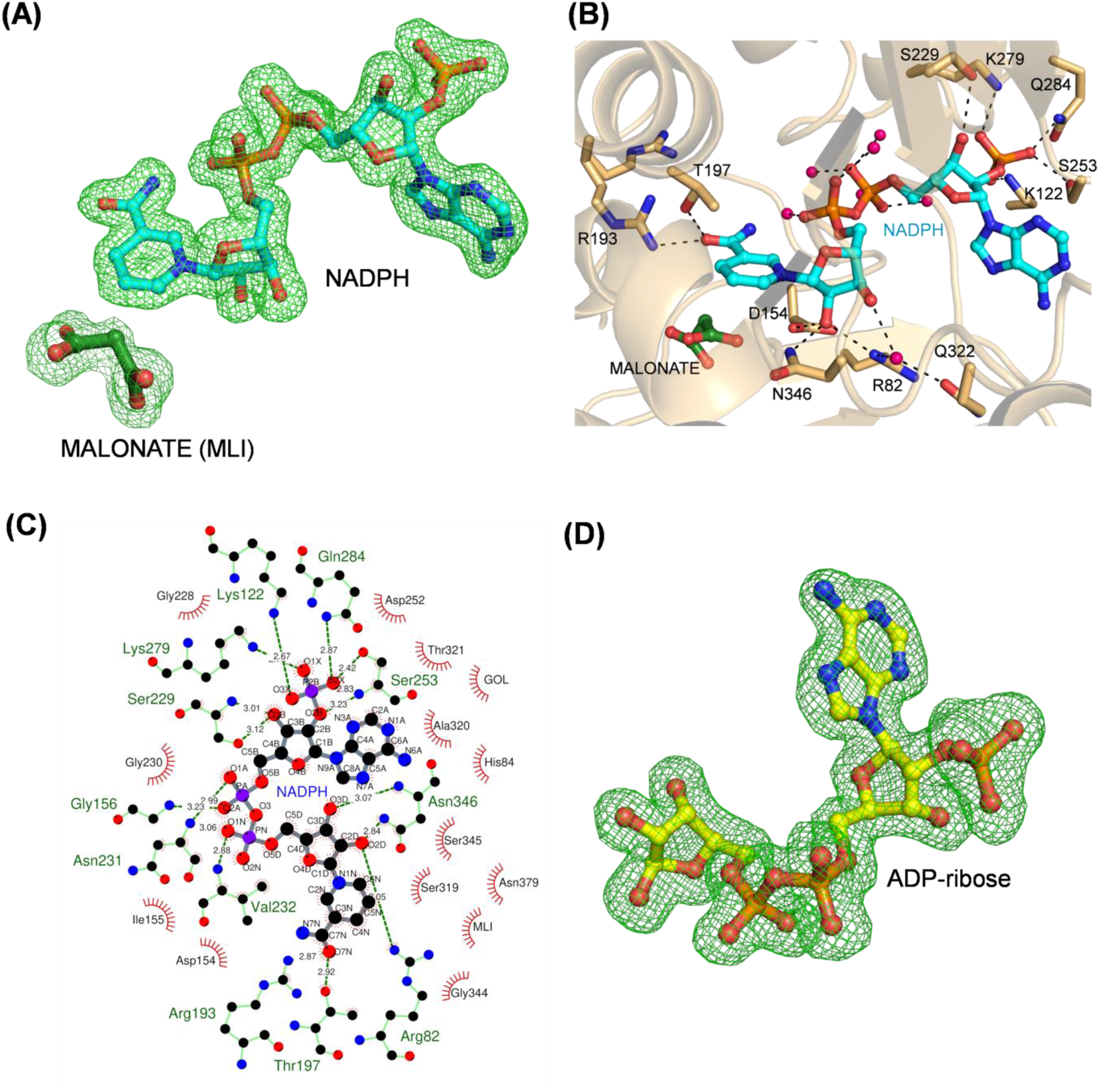
Binding of NADPH at the active site of AtGDH. (A) The sigma-A weighted *F*_o_-*F*_c_ omit map contoured at 3.0 σ level is shown as green mesh around refined NAPDH and malonate bound in the structure. (B) NADPH (cyan carbon) interacts with the active site residues (orange carbon) and the water molecules (pink sphere) *via* the hydrogen bonds (dashed lines). (C) Representation of interactions of NADPH (grey bond) with the active site residues (green bond) of AtGDH. The hydrogen bonds are shown as dashed green lines with distance in Å and the hydrophobic interactions as red semi-circle radiating lines. The carbon, oxygen and nitrogen are shown in black, red and blue color spheres, respectively. The figure was made using the LigPlot+ version 2.2 (3). (D) The presence of NADPH with cleaved nicotinamide ring (designated as ADP-ribose) in AtGDH-SIN-NADPH structure is identified by the sigma-A weighted *F*_o_-*F*_c_ omit map contoured at 3.0 σ level. Refined model of ADP-ribose is shown inside the map as ball and stick.

### Binding of the dicarboxylic acid metabolites at the active site of AtGDH

The substrate binding pocket in AtGDH lies inside the deep cavity formed by the conserved positively charged as well as neutral polar residues- Lys78, Gln99, Lys102, Lys114, Arg193, Asn379 and Ser386, as reported in other GDHs. The substrate binding pocket is mostly composed of the positively charged residues, therefore the metabolites bearing carboxylate group would have the general tendency to occupy this part of the active site. The presence of the substrate mimics-malonate, succinate and tartrate in the AtGDH crystal structure was confirmed by well-defined sigma-A weighted *F*_*o*_-*F*_*c*_ omit maps (Figures 4A, 5A and 5B). Their interactions with the active site residues are discussed below in details.

**Figure 5.**
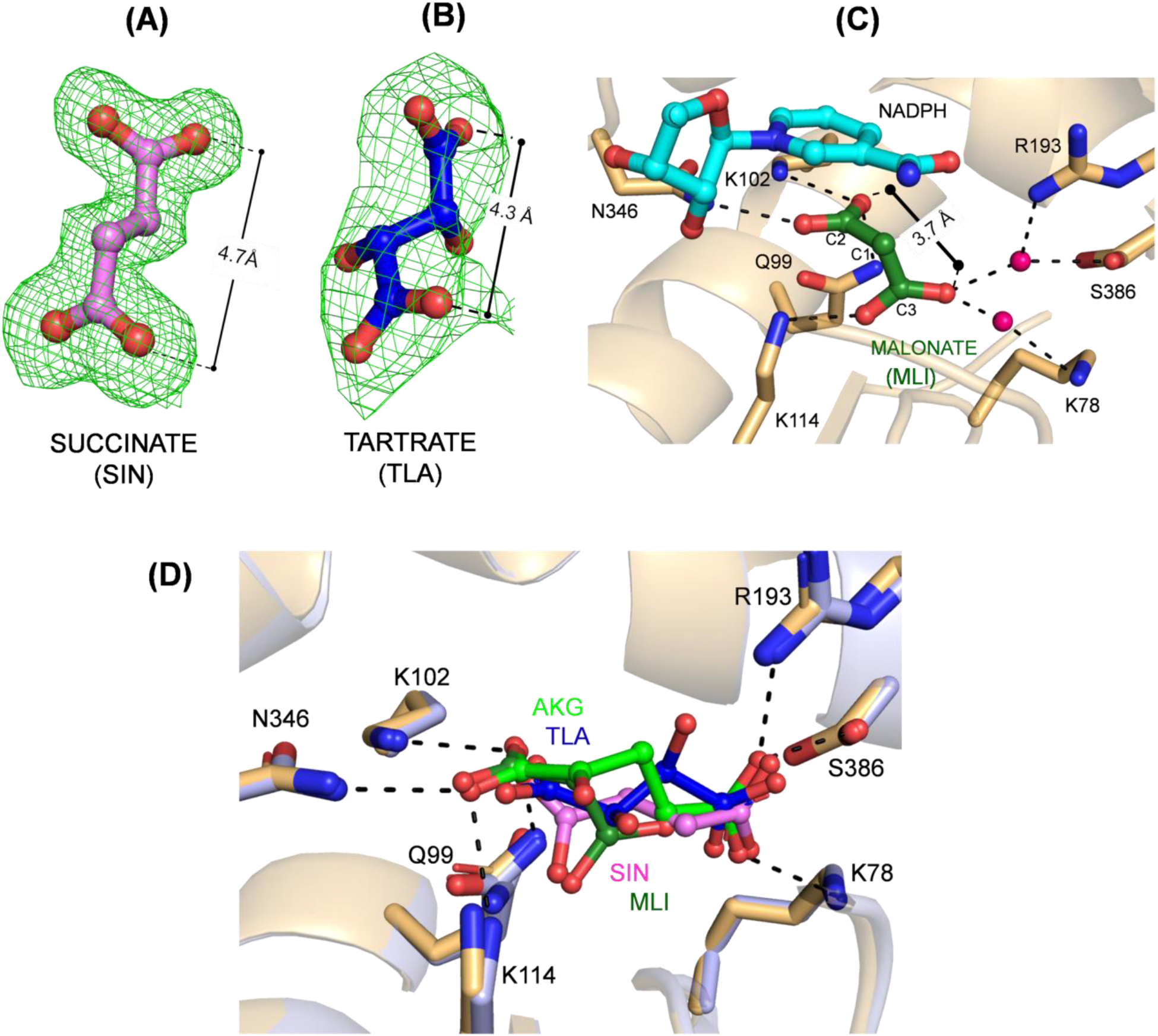
Binding and orientation of the substrate mimics-succinate, tartrate and malonate at the active site of AtGDH. The distance between the two carboxylate groups in a molecule is measured in Å. (A) The sigma-A weighted *F*_o_-*F*_c_ omit map contoured at 2.0 σ level is shown as green mesh around succinate (SIN; pink carbon) bound in AtGDH-SIN-NADPH structure and its inter-carboxylate distance is 4.7 Å. (B) The sigma-A weighted *F*_o_-*F*_c_ omit map contoured at 3.0 σ level is shown as green mesh around tartrate (TLA; blue carbon) bound in AtGDH-TLA-NADPH structure and its inter-carboxylate group distance is 4.3 Å. (C) Malonate with its inter-carboxylate group distance of 3.7 Å (MLI; dark green carbon) has polar interactions (dotted lines) with the active AtGDH site residues (orange carbon) and a water molecule (pink sphere).The carbon atoms of malonate are labeled as C1-C3. (D) Comparison of the active site pockets of substrate α-ketoglutarate (AKG) bound AnGDH ternary complex (5XVX, blue carbon) and dicarboxylate metabolite bound AtGDH (orange carbon) structures. The hydrogen bonding interactions for the substrate α-ketoglutarate (AKG; light green carbon) in AnGDH structure is shown as black dashed lines with the respective active site residues labeled.

A malonate molecule could be modelled in the well-resolved electron density observed in active sites of all the subunits in the AtGDH-MLI-NADPH structure (Figure 4A). In the solved structure it is observed that the substrate α-ketoglutarate which was used for making ternary complex for crystallization has been competitively replaced by malonate from the mother liquor. Malonate is a three carbon compound with two terminal carboxylate groups. The C2 carboxylate group forms hydrogen bond with the side chains of Gln99, Lys102, Lys114 and Asn346 (Figure 5C, S2A). It is also stabilized by a water mediated interaction with Asn379 backbone. The terminal C3 carboxylate group interacts with Lys78, Arg193 and Ser386 side chains *via* water molecules. The C3 carboxylate group also forms hydrogen bonds with the side chain Lys114 and Gly80 main chain. It is also stabilized by the water mediated interaction with Asn379 backbone. Malonate is oriented at the active site pocket such that the distance between the two carboxylates is 3.7 Å. Importantly, malonate being a small negatively charged molecule also binds at another site on the surface by interacting with side chain of Thr350, and the main chain of Gly327 and Glu328 in all three subunits. The carboxylate group(s) is also stabilised by the surrounding water molecules (Figure S2B).

The AtGDH-SIN-NADPH structure has well resolved electron density for succinate at the active site (Figure 5A). The α-ketoglutarate which was mixed with AtGDH prior to the setup for crystallization has been competitively replaced by succinate in the structure. Succinate is a four carbon compound with two terminal carboxylate groups; it stretches at the active site with an inter-carboxylate distance of 4.7 Å (Figure 5A). The terminal C1 carboxylate group of succinate forms hydrogen bonds with the side chains of Lys78, Arg193 and Ser386 residues while the C4 carboxylate group interacts with the Gly80, Gln99, Lys102 and Lys114 residues (Figure S2C).

The AtGDH-TLA-NADPH crystal diffracted to a lower resolution as compared to the other complexed crystals reported here. A noticeable electron density could be observed for L-tartrate molecule at the active site pocket (Figure 5B). Tartaric acid is a dibasic acid (4-carbon) naturally found in fruits like tamarind and grapes. It has two stereo-centers with two functional groups- the hydroxyl groups and terminal carboxylate groups. It orients with an inter-carboxylate group distance of 4.3 Å at active site pocket (Figure 5B). The C1 carboxylate group forms hydrogen bond with the Lys78, Arg193and Ser386 residues. The hydroxyl group at C2 position interacts with the side chain of Arg193 and the amide group of nicotinamide ring and at C3 position interacts with main chain of Gly153. The terminal C4 carboxylate group which lies adjacent to the nicotinamide ring of NADPH interacts with the side chain of Gln99, Lys102, and Lys114 residues (Figure S2D).

All the three ternary complex structures are nearly identical to each other. The dicarboxylates– malonate, succinate and tartrate occupy almost the same region of AtGDH active site by stabilizing interaction of varying hydrogen bond distances with the residues (Lys78, Lys114, Arg193 and Ser386) (Figure 5D, 6B). A structural comparison of AnGDH ternary complex (5XVX) with these three AtGDH complexes indicates that the three dicarboxylates occupy the α-ketoglutarate binding site of the enzyme (Figure 5D). As the AtGDH-SIN-NADPH structure has cleaved nicotinamide moiety, the Asn379 is more flexible (double conformation) and thus mediate interactions with succinate. Tartrate has maximum interactions and forms an additional hydrogen bond interaction with the Asp154, when compared to succinate and malonate. Notably, one subunit of the previously reported hexameric structure (5GUD) of *C. glutamicum* GDH (CgGDH) shows open conformation although it has citrate and NADP^+^ bound in the active site. A structural comparison of CgGDH with AtGDH-TLA-NADPH shows a slightly different binding mode of citrate as compared to tartrate (Figure S3). This could be due to the presence of an extra carboxylate in citrate. The AtGDH structures presented here provide sufficient direct evidences to suggest that the metabolites bearing carboxylate groups do bind to the active site of GDH and thereby cause enzyme inhibition.

**Figure 6.**
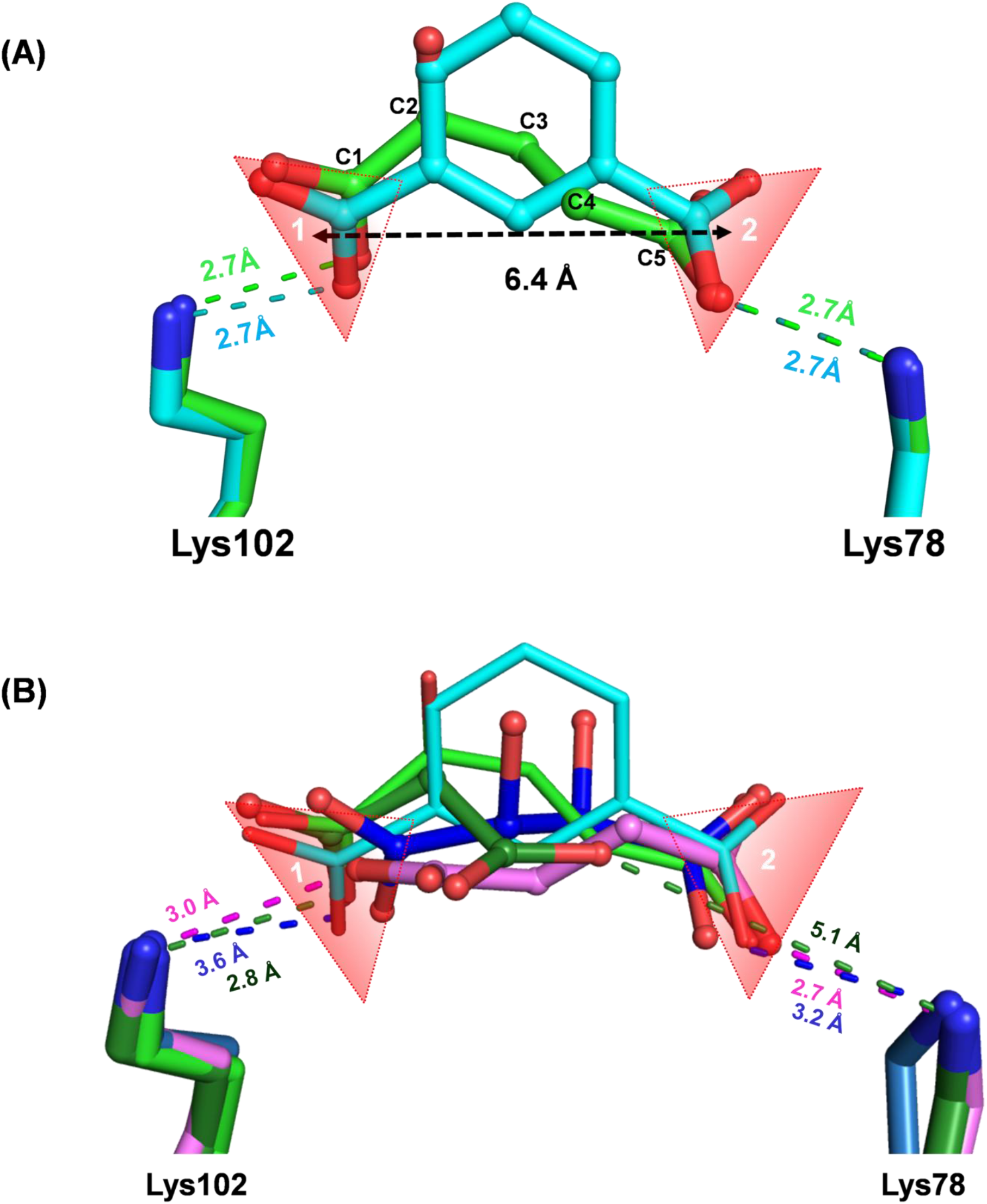
Molecular basis of inhibition of AtGDH by dicarboxylic acid metabolites. The inter-carboxylate distance (black dashed line) is measured in Å between the two carboxylate group (labelled as 1 and 2) in shaded red triangle. The electrostatic interactions of carboxylate groups with the conserved Lys78 and Lys102 residues in the active site and their corresponding distances (Å) are shown by dashed lines. (A) Superposition of the AnGDH structures bound to substrate-α-ketoglutarate (green ball and stick) (5XVX) and complexed with inhibitor isophthalate (cyan ball and stick) (5XW0). The C1–C5carbon atoms of α-ketoglutarate are labelled. (B) The AtGDH structures with bound substrate mimics-malonate (dark green carbon), succinate (pink carbon) and tartrate (blue carbon). Isophthalate (cyan stick) and α-ketoglutarate (light green stick) from superimposed AnGDH structures are also shown as reference.

## Discussion

Glutamate dehydrogenase (GDH) is an essential enzyme which interlinks the catabolic and biosynthetic pathways in all living systems. Despite numerous structural and biochemical studies in last 45 years several unique features of this enzyme are still not understood. GDH is well known to be regulated by a few metabolites arising from TCA cycle, glycolysis and others, in both allosteric and non-allosteric manner. We have expressed AtGDH as His-tagged recombinant protein in its active form. The enzymatic activity of AtGDH after removal of His_6_-tag is comparable to the same enzyme expressed without any affinity tag. We have determined the high resolution crystal structures of *A. terreus* GDH (AtGDH) with bound dicarboxylic acid metabolites (namely malonate, succinate and tartrate) as these compounds were present at a high concentration in the mother liquor solutions. Our biochemical data further demonstrate that AtGDH is indeed inhibited by dicarboxylic acid metabolites. Structural analysis provides the molecular basis of the mode of inhibition and explores the vulnerability of the active site of GDHs.

We initiated the crystallization experiments to obtain the AtGDH crystals as binary (NADPH bound) and ternary (AKG and NADPH bound) complexes. However, to our surprise, after solving the crystal structures we observed that AtGDH got crystallized with dicarboxylic acids from mother liquor along with NADPH. AtGDH complexes reported here do not have a full complement of substrates in the active site. Nevertheless, it is interesting that all these structures bound to NADPH (and the dicarboxylate metabolites) are in super closed conformation, similar to that observed for AnGDH ternary complex.^9^ Surprisingly, succinate and malonate individually, from the mother liquor could replace α-ketoglutarate from the active site; as the AtGDH-SIN-NADPH and the AtGDH-MLI-NADPH structures were determined from the crystals grown using the enzyme that was pre-incubated with the substrates – α-ketoglutarate and NADPH. The architecture of active site of GDHs from different origins is well conserved and aligned by the positively charged residues which facilitate binding of the negatively charged carboxylate group containing molecules. It is important to note that the super closed conformation is usually triggered by both substrate and coenzyme binding, but here we present the first super closed conformation of GDH structure captured without the substrate α-ketoglutarate. These AtGDH complexed structures indicate that the binding of the dicarboxylate group containing metabolites along with the coenzyme could induce catalytically competent closed conformation in GDHs. On the contrary, the open conformation in citrate and NADP^+^ bound CgGDH structure (5GUD) might be due to the extra carboxylate group present in citrate.^47^

Several studies state the abnormal accumulation of organic metabolites during the rare metabolic disorder organic aciduria.^25^ Our biochemical studies show that malonate, maleate, succinate and fumarate inhibit the forward reaction of AtGDH almost to the same extent (~30– 40% decrease in residual activity at 20 mM) while the tartrate barely inhibits at 20 mM concentration. Although tartrate is stabilised by the more number of polar interactions with the surrounding residues as compared to succinate and malonate, it is a very poor inhibitor (<30% inhibition at 50 mM). Comparatively, malonate exhibits the maximum inhibitory potency over fumarate, maleate, succinate and tartrate. The crystal structure comparison of AnGDH (5XW0, 5XVX) with the AtGDH structures reported in this study illuminate the possible molecular basis of GDH inhibition by dicarboxylic acid containing compounds. Earlier Caughey et al (1957), albeit without structural data, reported that the metabolites with planar conformation and a terminal inter-protonic distance of ~7.65 Å can act as good inhibitors for GDH.^28^ The known potent inhibitors of GDH such as isophthalate, 2-methyleneglutarate, oxalylglycine and glutarate might orient and stretch in a way to generate a 5-carbon molecule representation^9,28,36^ when they get bound in the enzyme active site. The recent crystal structures of AnGDH reveal that the carboxylate groups of α-ketoglutarate and isophthalate occupy the same positions with separation of ~6.4 Å (Figure 6A). The bound α-ketoglutarate does not adopt a planar and maximum extended conformation in the catalytically competent active site of AnGDH. Interestingly the carbon atoms of the substrate and inhibitor occupy different three dimensional space. It appears that the spatial separation of the two carboxylate groups and their binding (by salt bridge interactions) to two conserved lysine residues (Lys78 and Lys102 in AnGDH) in the GDH active site are the important factor that dictate the inhibitory potential of the inhibitors. The compounds with the carboxylate groups separated by ~6.4 Å and make salt bridge interactions with both the lysine residues would be highly potent inhibitors (e.g. isophthalate). Notably, the substrate mimics – malonate, succinate and tartrate as dicarboxylate anions occupy the region of AtGDH binding site very similar to that of α-ketoglutarate in the AnGDH active site. We hypothesize that fumarate and maleate being structurally similar to succinate could possibly occupy the same active site as well. However, certain differences in their binding mode might be the reasons why they act as poor inhibitors. Malonate has the highest inhibitory potency among other substrate mimics, and has one of its terminal carboxylate groups occupying almost the identical position (near Lys102) as that of the C1 carboxylate of α-ketoglutarate. The succinate has one of its terminal carboxylate groups occupying the same position (near Lys78) as that of the second carboxylate (C5) of α-ketoglutarate (Figure 6B). While the second carboxylate of succinate does not occupy the position (near Lys102) as that of the C1 carboxylate of α-ketoglutarate (Figure 6A, 6B). It is evident that succinate or malonate forms only one strong salt bridge interaction with the nearby lysine residues (Lys102 or Lys78). The inability of malonate or succinate to mimic the exact salt bridge mediated binding pattern of α-ketoglutarate makes them weaker inhibitors as compared to isophthalate. However, slight higher inhibitory potency of malonate as compared to succinate might be due to its interaction with Lys102 which allow the former compound to orient somewhat similar manner as substrate in the active site. Notably, although tartrate is a 4-carbon containing compound with maximum active site interactions, have its carboxylate groups shifted slightly away from the actual binding pocket due to presence of two hydroxyl groups. This difference in binding mode could account for the low inhibitory potency of tartrate. Interestingly, maleic acid which accumulates during PAcidemia condition is reported to inhibit human kidney GDH activity.^22^ The inhibition potency of maleic acid might be related to better positioning of the two carboxylate groups in the mammalian GDH active site due to presence of the *cis* double bond between C2 and C3. However, the fumarate (*trans*) and maleate (*cis*) act as weak inhibitors for AtGDH as compared to mammalian GDH. The inhibitory action of these dicarboxylic acid metabolites might vary among GDHs from different sources. It could be due to the variation among the surrounding residues at the active site pocket which dictate their binding affinity and inhibitory potency. Structural comparison indicates presence of Met115 in human GDH (PDB ID: 1NR1) and Met111 in bovine GDH (PDB ID: 3JD1) near the binding pocket of substrate, whereas Gln99 occupies the identical position in AtGDH. Furthermore, the experimental *in-vivo* and structural data as well as more detailed biochemical studies might explain the reason behind the differential inhibitory effects of these dicarboxylic acid metabolites. Our structural analysis points out that Cα substituted 5-carbon dicarboxylate-containing molecules with the separation of the two carboxylate groups by 6.4 Å would be potent inhibitors over 3-, 4- or 5-carbon dicarboxylic molecules against GDH activity. Therefore, the poor inhibition of AnGDH by 3,5-pyrazoledicarboxylate and Δ^1^-piperidine-2,4-dicarboxylate as compared isophathalate^2^ might be due to the shorter spatial distance between the carboxylate groups of former two compounds as compared to the third. The results presented in this study provide a plausible rationale for the inhibition of the GDH activity by dicarboxylate metabolites, and would aid in understanding the mechanism of binding of other potent inhibitors of GDHs.

## Supporting Information

The supporting information can be found at the end of this article

## Acknowledgements

We acknowledge the support from Protein Crystallography Facility at IIT Bombay for the protein crystallization and data collection. We are thankful to Dr. Nupur Agarwal for her guidance on the enzyme assays. The work was supported by Ramalingaswami Re-entry Fellowship (Department of Biotechnology, Ministry of Science and Technology, India) and research seed grant from IRCC, IIT Bombay (to P. B.). This work was partially supported by a research grant [37(1)/14/04/2017-BRNS/37040] to N. S. P. from Board of Research in Nuclear Sciences, DAE, Mumbai, India. Fellowship to B. K. J. G. from Department of Biotechnology (DBT), Ministry of Science and Technology, India is also acknowledged.

## Data availability

The coordinates and structure factors for the AtGDH complexed structures are deposited in the Protein Data Bank (PDB) (http://www.rcsb.org) with the accession codes 7ECR, 7ECS and 7ECT.

## Supporting Information

**Supplementary Figure 1.**
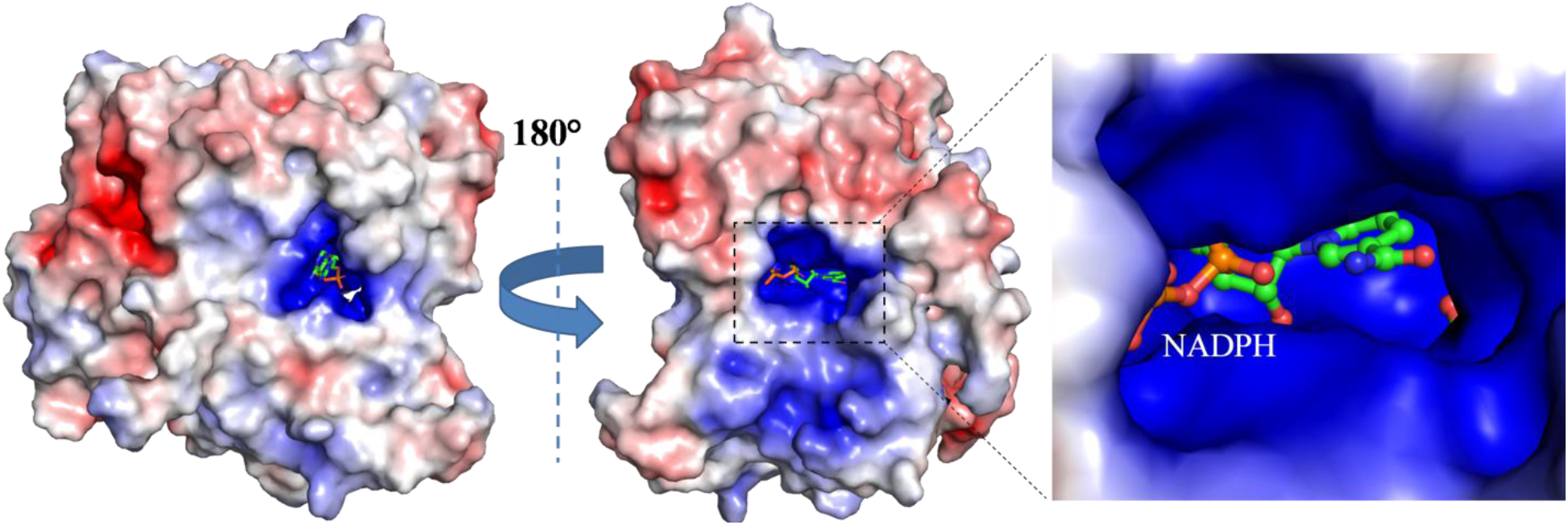
Electrostatic surface charge distribution on AtGDH (left) and after 180° rotations around the vertical axis (right). The inset shows zoomed in view of the strongly positively charged binding pocket with coenzyme NADPH and malonate bound. Color code of the charged surface; red: negative, blue: positive and white: neutral. The figure is generated by the APBS tool of PyMol (4).

**Supplementary Figure S2.**
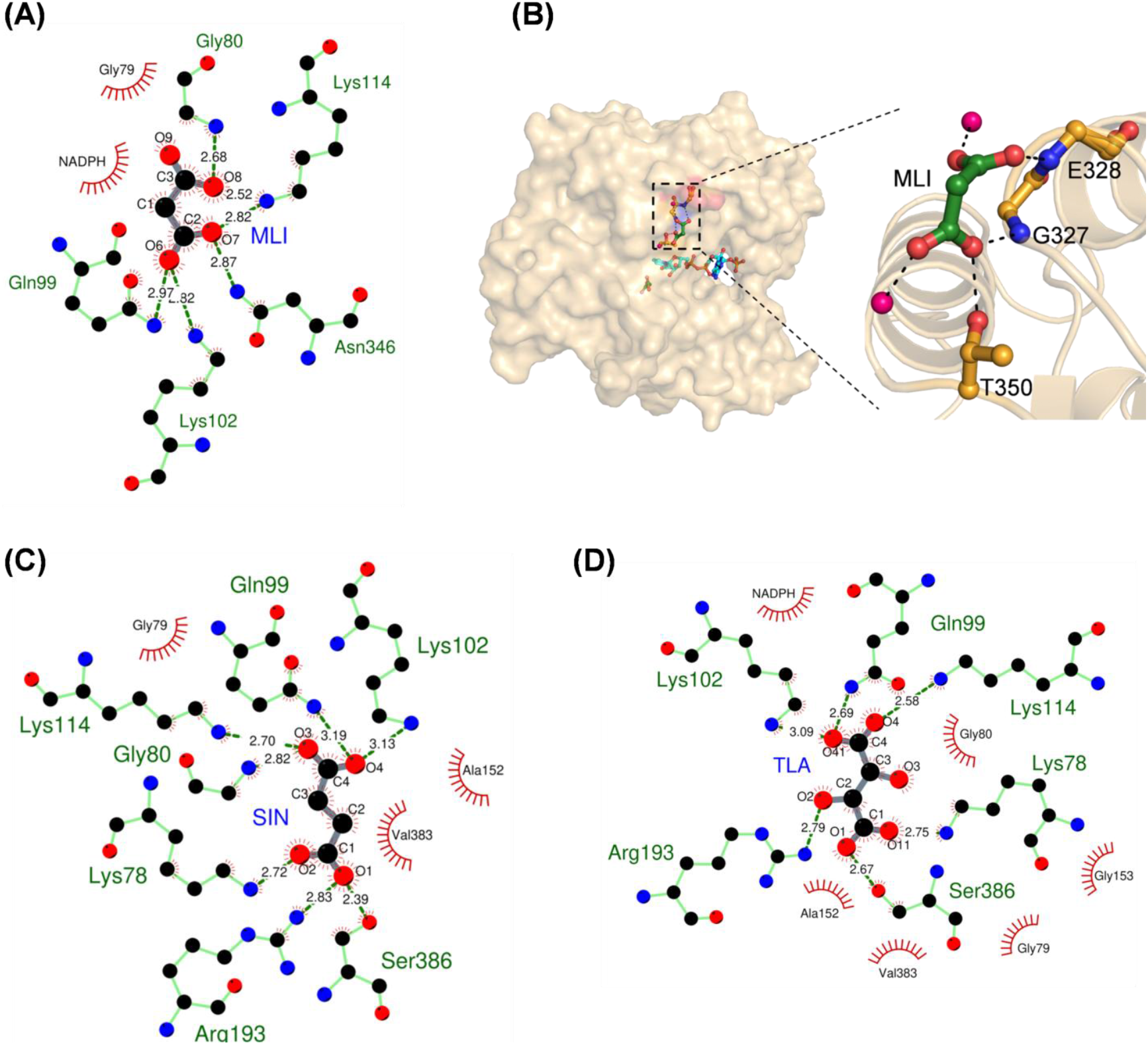
Interactions of substrate mimics. Schematic representation of dicarboxylate metabolites hydrogen bond interactions at the active site of AtGDH analzed by LigPlot (3).The hydrogen bond interactions are shown as dashed green lines with distance in Å and the hydrophobic interactions as red semi-circle radiating lines in the AtGDH active site (green bond) with (A) malonate (MLI), (C) succinate (SIN) and (D) tartrate (TLA). The carbon, oxygen and nitrogen are shown in black, red and blue color sphere, respectively. (B) Surface representation of AtGDH (orange) with the bound malonate (dark green stick) on another binding pocket away from the active site. The inset represents the zoom in view of the hydrogen bond interactions (dashed lines) formed by malonate (green carbon) with the surface residues (orange carbon) and two water molecules (pink spheres).

**Supplementary Figure 3.**
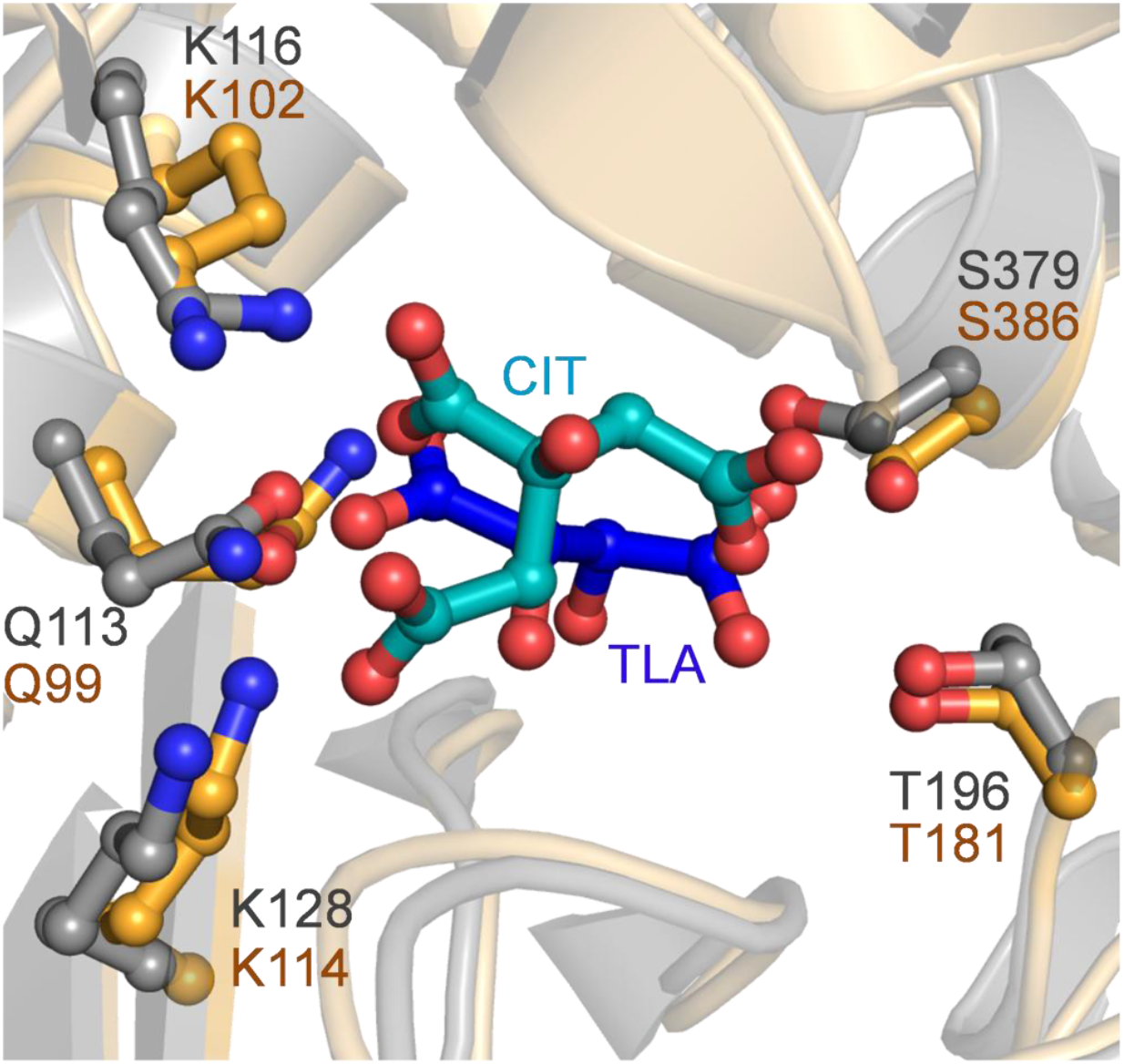
Differences in the mode of binding of citrate and tartrate at the active site of GDHs. The superposition of the *C. glutamicum* GDH structure (5GUD) (grey carbon) complexed with citrate (labeled as CIT; deep cyan carbon) and AtGDH-TLA-NADPH (orange carbon) with bound tartrate (labeled as TLA; dark blue carbon) at the active site. The active site residues are shown in colors corresponding to their proteins.

